# Hijacking and recording intracellular RNAs in human cells using eukaryotic reprogrammed tracrRNAs

**DOI:** 10.1101/2024.10.28.620760

**Authors:** Seok-Hoon Lee, Junho Park, Chanju Jung, Sangsu Bae

**Author notes:** To whom correspondence should be addressed to S.B. These authors contributed equally to this work (Seok-Hoon Lee, Junho Park).

## Abstract

Recording intracellular RNA expression events is critical for understanding cell states and biological processes, but no method without responsive promoters is available in mammalian cells. Here, we determined the sequence formula of eukaryotic reprogrammed tracrRNA, named euRptr, which can hijack intracellular RNAs to make a complex with CRISPR-associated tools. By leveraging the programmability of euRptr, we devised the system “CHEETAH” to directly capture and record intracellular RNAs in human cells. We validated that CHEETAH induces self-editing by hijacking endogenous mRNAs, that CHEETAH selectively records different viral RNA fragments on a DNA reporter using base editors with a shared-euRptr, and that CHEETAH enables multiplexed, time-resolved recording of viral RNAs on a DNA Tape using prime editors, providing a new avenue for investigating intracellular RNA in eukaryotes.

## Introduction

Cellular behavior is governed by complicated biological processes that occur within living systems and understanding how molecular changes unfold over time is essential to elucidating the causes and progression of various biological phenomena. The CRISPR-Cas (Clustered Regularly Interspaced Short Palindromic Repeats and CRISPR-associated) system, a bacterial adaptive immune system (*1-8*), has been harnessed to record past molecular events by creating modifications in DNA, a stable substrate as a memory device (*9-19*). Initial recording techniques utilized signal-responsive promoters expressing Cas protein and single guide RNA (sgRNA), an engineered combination of CRISPR RNA (crRNA) and trans-activating crRNA (tracrRNA) (*9-13*). However, these approaches are constrained by the diversity and number of input signals that they can process. For instance, infections by multiple distinct viruses are challenging to record in a multiplexed or virus-specific manner using the signal-responsive promoter.

Alternatively, new approaches that directly capture DNA molecules using Cas1-Cas2 (*14-16*) or that directly capture RNA molecules using RT-Cas1-Cas2 (*17, 18*) and using reprogrammed tracrRNA (Rptr) (*19, 20*) have been developed and applied in bacteria. However, it remains unclear whether these methods are effective in mammalian cells, in which the intracellular environment is more complicated by a membrane-bound nucleus and the presence of chromatin compared to the intracellular environment of bacterial cells.

Herein, we identified consensus sequences for eukaryotic reprogrammed tracrRNA (termed euRptr) with Cas9 nucleases in human cells and successfully developed CHEETAH (CRISPR-based Hijacking of Endogenous and Exogenous Transcripts for Annotation in Human cells), a system for mammalian cells. We demonstrated that the CHEETAH could hijack endogenous messenger RNAs (mRNAs) and result in generation of insertion and deletion (indel) mutations at the encoding genes in human cells, indicating an euRptr-leveraged self-editing. Moreover, we demonstrated that the CHEETAH could implement RNA recording of transfected viral RNAs within the encoding DNA by means of base editor (BE) (*21, 22*) and prime editor (PE) (*23*) platforms in human cells, indicating an euRptr-leveraged RNA recording on DNA.

## Results

### Determination of euRptr formula for NG-SpCas9–mediated gene editing in human cells

CRISPR-Cas nucleases typically display sufficient activity in prokaryotic cells, but many of them exhibit poor activity in eukaryotic cells due to more complicated environment of eukaryotic cells (*24-28*). Although the previous cases demonstrated the hijacking of bacterial RNAs with Cas9 from *Campylobacter jejuni* NCTC11168 (CjCas9) *in vitro* (*20*) and *Streptococcus thermophilus* (Sth1Cas9) in *Escherichia coli* (*19*), Cas9 from *Streptococcus pyogenes* (SpCas9) exhibits the highest editing activity in mammalian cells and there are several engineered variants of SpCas9 with expanded protospacer-adjacent motif (PAM) sequences (*29-31*). Therefore, we used SpCas9 to explore the potential of hijacking cellular RNAs in human cells. For the crRNA:tracrRNA hybridization, we sought to identify eukaryote-specific rules to reprogram the tracrRNA anti-repeat sequence in human cells (**Fig. 1A**). The determination of the consensus repeat sequences is crucial to reprogram the tracrRNA and hijack cellular RNAs with the target motifs.

**Figure 1.**
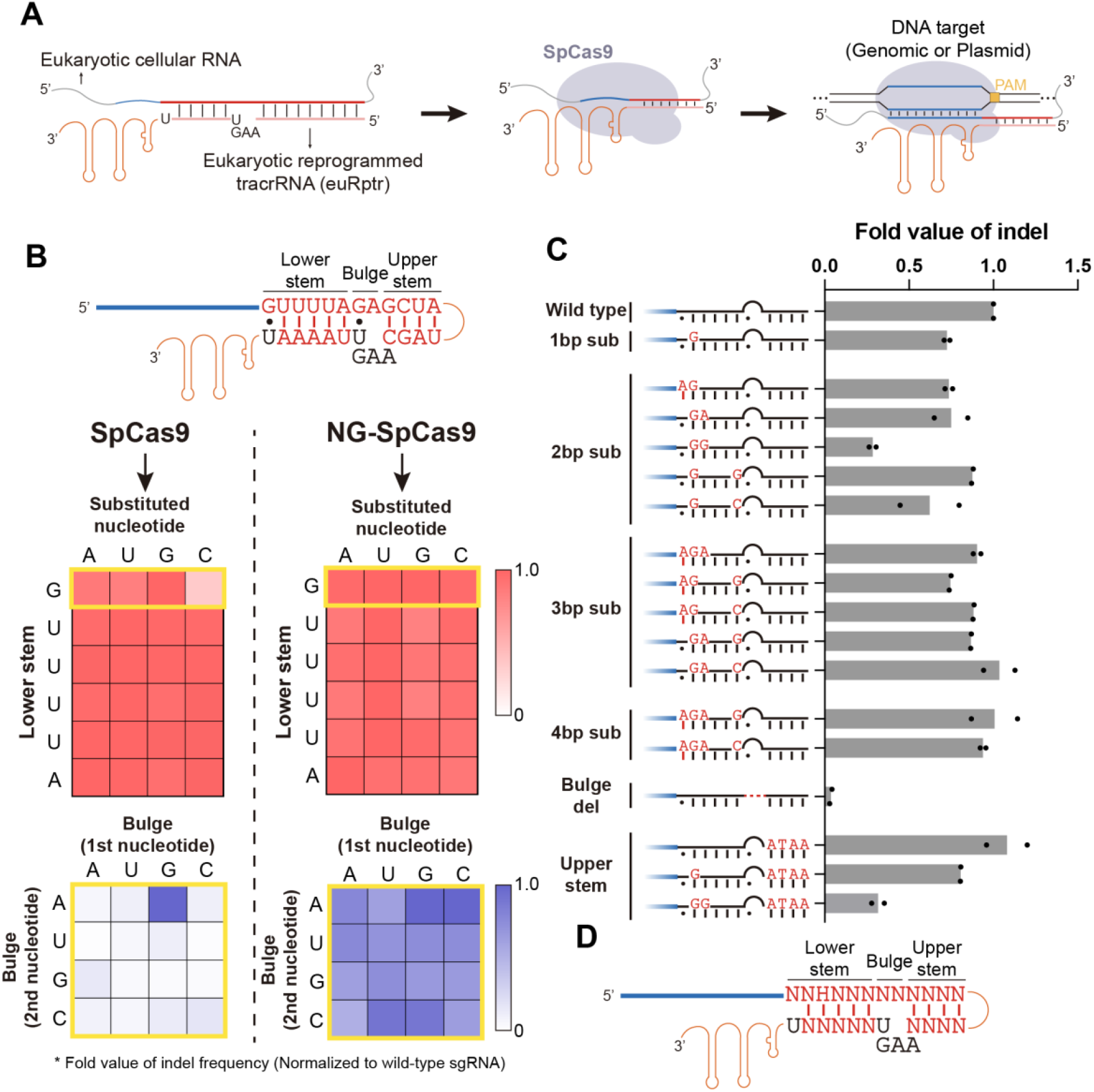
Evaluation of sgRNA scaffold tolerance for SpCas9 and NG-SpCas9. **(A)** Schematic of the hijacking system (CHEETAH) in human cells. The blue and red regions within cellular RNA indicate the spacer and hijacking target sequences, respectively. The pink region marks the reprogramming area within the tracrRNA. The sequences within the secondary structure of tracrRNA were fixed to minimize disruption of editing efficacy. **(B)** The effect of simple substitutions on the lower stem and bulge regions of *TBP* gene-targeting sgRNA. The nucleotide notation in the heatmap indicates the nucleotide in the repeat sequence. In the lower stem (5’-GUUUUA-3’), each nucleotide was independently substituted with all other nucleotides. In the bulge (5’-GA-3’), every combination of substitutions was tested. All heatmap values represent the fold change relative to unmodified sgRNA (*n* = 3). Yellow boxes indicate positions related to RNA secondary structure. **(C)** The effect of complex mutations on the lower stem, bulge, and upper stem. Mutations are indicated in red. The upper stem mutations (ATAA) correspond to the same positioned sequence in the *TBP* target site. For base-pairing annotation, dots indicate Wobble base pairs, and dash lines indicate Watson-Crick base pairs. All values represent the fold change compared to unmodified sgRNA (*n* = 2). **(D)** Summary of the reprogrammable sequences in the repeat:anti-repeat region within sgRNA. The “H” in the lower stem is required to avoid the 5’-NGG-3’ mutation, which is a disruptive mutation as shown in Figure 1C. This rule is used to choose the hijacking target region of the RNA of interest and design reprogrammed tracrRNAs. N indicates A, T, G, or C, and H indicates A, T, or C.

To examine the tolerable sequences of the crRNA:tracrRNA hybrid, we targeted the *TBP* gene and prepared plasmids expressing a sgRNA with one or more mutations within the lower stem, bulge, and upper stem regions of the sgRNA scaffold. The RNA secondary structure related-sequences within the anti-repeat sequences, including the 3’-U-5’ of G-U wobble base pair and the 3’-UGAA-5’ of the bulge, were fixed to minimize disruption of editing efficacy. After SpCas9 and each sgRNA variant were transfected into HEK293 cells, gene editing efficiencies at the *TBP* site were measured by targeted deep sequencing. Tolerance to variation was lowest in the bulge region and the G-U wobble base pair of the lower stem of SpCas9 (**Fig. 1B** and **Fig. S1**), indicating that SpCas9 might not be suitable for reprogramming the tracrRNA. Thus, we investigated the tolerance of the sgRNA scaffold with NG-SpCas9 that recognizes 5’-NG-3’ sequences as a PAM and offers a broader range of target sites than the canonical SpCas9 (*29*). NG-SpCas9 tolerated single substitutions in most scaffold regions (**Fig. 1B**) and accepted more complex mutations, except in the 5’-NGGNNN-3’ sequence of the lower stem or whole deletion of the bulge region (**Fig. 1C** and **Fig. S1**). When we used adenine base editor (ABE) instead of Cas9 nucleases, we observed a similar tendency: ABE8e recognizing NGG PAM had lower tolerance to variation than NG-ABE8e recognizing NG PAM (**Fig. S2**) (*32*). These results suggested that reprogramming of tracrRNA with NG-SpCas9 or NG-ABE8e has a potential to hijack intracellular RNAs with a wide targeting scope in human cells. Through systematic mutation tolerance analysis, we determined 5’-NNHNNNNNNNNN-3’ for sequences of lower stem to upper stem as optimal for using euRptr with NG-SpCas9 or NG-ABE8e and cellular RNA in human cells (**Fig. 1D**).

### euRptr hijacks endogenous mRNAs and induces gene editing in human cells

We hypothesized that when NG-SpCas9 is transfected with a designed euRptr into human cells, the NG-SpCas9::euRptr::hijacked mRNA ribonucleoprotein (RNP) complex will form and co-transport into the nucleus due to the nuclear localization signal (NLS) in the Cas protein. Consequently, self-editing at the endogenous mRNA-expressing gene will occur if the target contains a suitable PAM sequence, 5’-NG-3’ (**Fig. 2A**). The criteria for designing the euRptr target are as follows: (i) the euRptr target sequences must be located in exon regions because mature mRNA must contain the euRptr-corresponding sequences after the RNA splicing process, (ii) the euRptr target must involve the following consensus sequence, 5’-NGHNNNNN-3’ for the sequences of lower stem to bulge, to hybridize with the designed euRptr and induce gene editing through NG-SpCas9, and (iii) to functionally substitute for the cRNA, the mRNAs should be abundantly transcribed in cells. Although the anti-repeat sequence includes the complement consensus sequence of 3’-UCDNNNUGAA-5’ at the sequence from the lower stem to bulge, we used two different versions of euRptr: (i) euRptr-v1 that contains a 33-nt anti-repeat sequence, same as the length of original tracrRNA; (ii) euRptr-v1s that contains a short 22-nt anti-repeat sequence, corresponding to the length of the trimmed tracrRNA during crRNA maturation by RNase III in CRISPR-Cas system.

**Figure 2.**
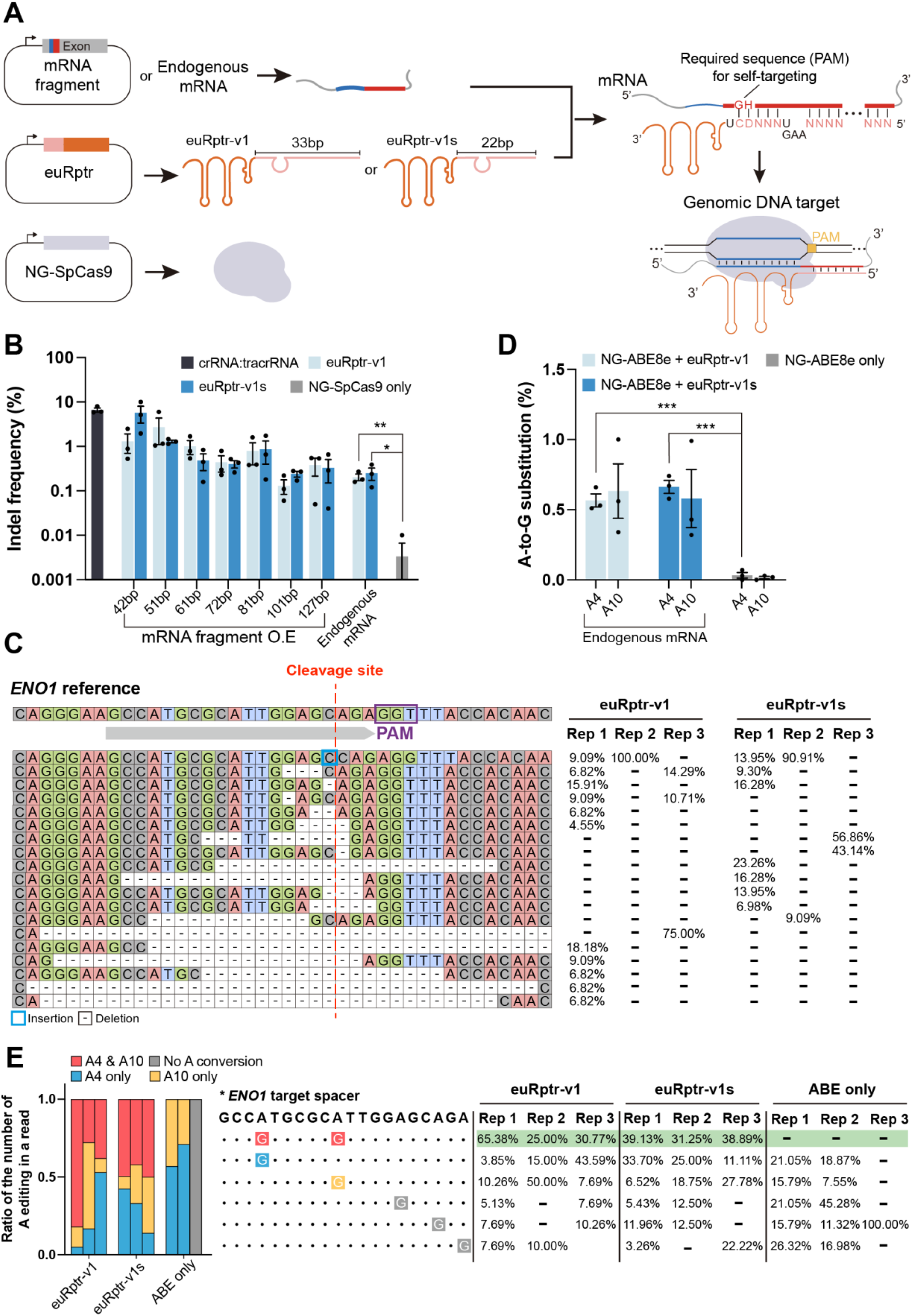
Genomic self-targeting with euRptr recognizing an overexpressed mRNA fragment or endogenous mRNA. **(A)** Schematic representation of the experiments in the figure. Both euRptr and NG-SpCas9 were overexpressed from plasmids. The RNA of interest was either an overexpressed mRNA fragment or endogenous mRNA, depending on the experiment. For self-targeting, the rule (5’-NGHNNN-3’ downstream of the spacer) was applied to search for endogenous targets. The sequence should be in the exon, and the 5’-NGH-3’ sequence acts as a PAM for the self-targeting. **(B)** Indel frequency at the *ENO1* endogenous target site. Overexpressed *ENO1* mRNA fragments or endogenous *ENO1* mRNA were used with euRptr and NG-SpCas9 (*n* = 3; error bars represent mean ± SEM). *P* = 0.0042 for euRptr-v1 and *P* = 0.0343 for euRptr-v1s. **(C)** Indel patterns generated by hijacking endogenous *ENO1* mRNA. The percent ratio of each editing pattern within the entire set of mutations is described in the right panel of this figure. The overall indel frequencies including the wild-type pattern are 0.27%, 0.18%, and 0.16% for euRptr-v1 and 0.24%, 0.12%, and 0.39% for euRptr-v1s. **(D)** A-to-G substitution rate at the *ENO1* endogenous target site. Endogenous *ENO1* mRNA was used with euRptr and NG-ABE8e (*n* = 3; error bars represent mean ± SEM). *P* = 0.0005 for euRptr-v1 and *P* = 0.0002 for euRptr-v1s. **(E)** Base-editing patterns generated by hijacking endogenous *ENO1* mRNA. A unique bystander editing pattern was observed in the experimental group but not in the control group (Green box). The described numbers indicate the percent ratio of each editing pattern within the entire mutations. The overall base editing frequencies, including wild-type pattern, are (A4; 0.51%, A10; 0.56%), (A4; 0.53%, A10; 1.00%), and (A4; 0.66%, A10; 0.34%) for euRptr-v1, (A4; 0.67%, A10; 0.43%), (A4; 0.58%, A10; 0.32%), and (A4; 0.74%, A10; 0.99%) for euRptr-v1s, and (A4; 0.03%, A10; 0.03%), (A4; 0.07%, A10; 0.02%), and (A4; 0.00%, A10; 0.00%) for the control group. *P*-values in B, D were calculated using Student’s *t*-test (* *P* < 0.05, ** *P* < 0.01, *** *P* < 0.001).

Based on these criteria, we selected *ENO1, GAPDH*, and *RPS17* genes that are abundantly expressed in HEK293 cells (**Fig. S3A**) and searched for suitable target sites containing the formula of 5’-NGHNNNNN-3’. For the *ENO1* gene, a target site containing 5’-GGUUUACC-3’ was selected. As a positive control, we tested a set of canonical crRNA that contains 5’-GUUUUAGA-3’ at the lower stem to bulge region and tracrRNA that contains 3’-UAAAAUUGAA -5’ at the lower stem to bulge region with NG-SpCas9. Targeted deep sequencing at the endogenous site showed an indel frequency of approximately 6.63% (**Fig. 2B**). We next tested CHEETAH by introducing NG-SpCas9 and *ENO1* target-specific euRptr that contains 3’-UCAAAUUGAA-5’ at the lower stem to bulge region together with a plasmid overexpressing each *ENO1* mRNA fragment containing 5’-GGUUUACC-3’, ranged from 42 to 127 bp. Compared with the positive control (crRNA:tracrRNA), we observed lower but confident editing efficiencies for all cases: We detected approximately 1.30% (euRptr-v1) and 5.72% (euRptr-v1s) for the 42-bp mRNA fragment and 0.38% (euRptr-v1) and 0.33% (euRptr-v1s) for the 127-bp mRNA fragment (**Fig. 2B** and **S3B**). These results indicated that the overexpressed mRNA fragments served the same function as the crRNA, supporting the hypothesis that CHEETAH can induce euRptr-leveraged self-editing by hijacking cellular mRNAs without input crRNAs in human cells. Thus, we transfected only NG-SpCas9 and the *ENO1* target-specific euRptr into HEK293 cells and detected editing efficiencies of approximately 0.20% (*P* = 0.0042) and 0.25% (*P* = 0.0343) with euRptr-v1 and euRptr-v1s, respectively, at the target site in the *ENO1* gene (**Fig. 2B** and **S3B**). Furthermore, the editing outcomes exhibited typical indel patterns associated with Cas9-mediated gene editing (**Fig. 2C**) (*33*), supporting that CHEETAH enables self-cleavage and gene editing by means of the cellular *ENO1* transcripts. To further validate CHEETAH-mediated self-editing, we tested NG-ABE8e with the *ENO1* target-specific euRptr without any overexpression of mRNA fragments in HEK293 cells. Bystander adenines (*34, 35*), such as the fourth adenine (A4) and tenth adenine (A10), were simultaneously converted (**Fig. 2D** and **2E**), with the highest A-to-G conversion rates in A10 (1% for euRptr-v1 and 0.99% for euRptr-v1s) (**Fig. 2D**). Despite lower editing efficiencies that we observed with CHEETAH targeting *ENO1*, we also detected CHEETAH-mediated gene editing or base editing with NG-SpCas9 or NG-ABE8e, respectively, in *GAPDH* and *RPS17* by means of their endogenous mRNAs (**Fig. S4**). Taken together, we concluded that CHEETAH hijacks cellular mRNAs and induces self-targeting resulting in gene editing or base editing when combined with NG-SpCas9 or NG-ABE8e, respectively, in human cells.

### CHEETAH selectively records the existence of similar RNA fragments using base editors

We next investigated whether CHEETAH can detect intracellular expression of viral RNAs by hijacking the RNA fragments and recording the existence events on DNA. As a model system, we selected a SARS-CoV-2 open reading frame (ORF) 10 transcript (117 bp in total length) as an exogenous RNA because ORF10 is present in all sub-genomic RNAs of the SARS-CoV-2 virus (*36*). Contrary with the above experiments that aim to induce self-targeting to the mRNA-expressing gene, this application uses a DNA plasmid (pRecord) as a memory system into which CHEETAH records an existence event by hijacking the viral RNA. To maximize editing of pRecord, we selected target regions with the formula 5’-NHHNNNNN-3’ (sequence of lower stem to bulge) in which targets containing NG PAM were excluded to avoid self-targeting. We also prepared a target-specific pRecord corresponding to the selected euRptr. In this application, pRecord contains NG PAM. For the euRptr-leveraged RNA recording, we employed NG-ABE8e to write the presence of exogenous RNA transcripts on pRecord (**Fig. 3A**). We designed three euRptr candidates within the SARS-CoV-2 ORF10 transcript and prepared corresponding pRecord (pRecord-ORF10) for each euRptr. Resultantly, we found that both euRptr-v1 and euRptr-v1s showed moderate recording efficiencies for target 1 (average 8.5%) and target 3 (average 5.1%) (**Fig. 3B**), and that the RNA recording events on pRecord-ORF10 were maintained for at least 7 days (**Fig. S5**). However, we measured very low RNA recording rates (average 0.4%) for target 2 on pRecord-ORF10 with NG-ABE8e (**Fig. 3B**) and NG-SpCas9 (**Fig. S6A**), suggesting that the hybridization of euRptr::viral RNA fragment is sequence-dependent and inhibited by the RNA secondary structure of euRptr-v1 and euRptr-v1s. Therefore, we tested a different euRptr architecture (euRptr-v2 and euRptr-v2s) with a t-lock scaffold (*37*) forming a super-stable hairpin that prevents misfolding of the euRptr::viral RNA fragment. Both euRptr-v2 and euRptr-v2s resulted in enhanced RNA recording efficiencies for target 2, producing six or seven times the recording events compared to euRptr-v1 and euRptr-v1s (**Fig. 3B** and **Fig. S6B**), consistent with RNA secondary structure impacting the euRptr-leveraged hijacking process. Furthermore, RNA secondary structure prediction analysis also indicated higher stability of euRptr-v2s for target 2 (**Fig. S7**). Because we observed that euRptr-v2s generally outperformed euRptr-v2 (**Fig. 3C**), we used euRptr-v2s for further experiments.

**Figure 3.**
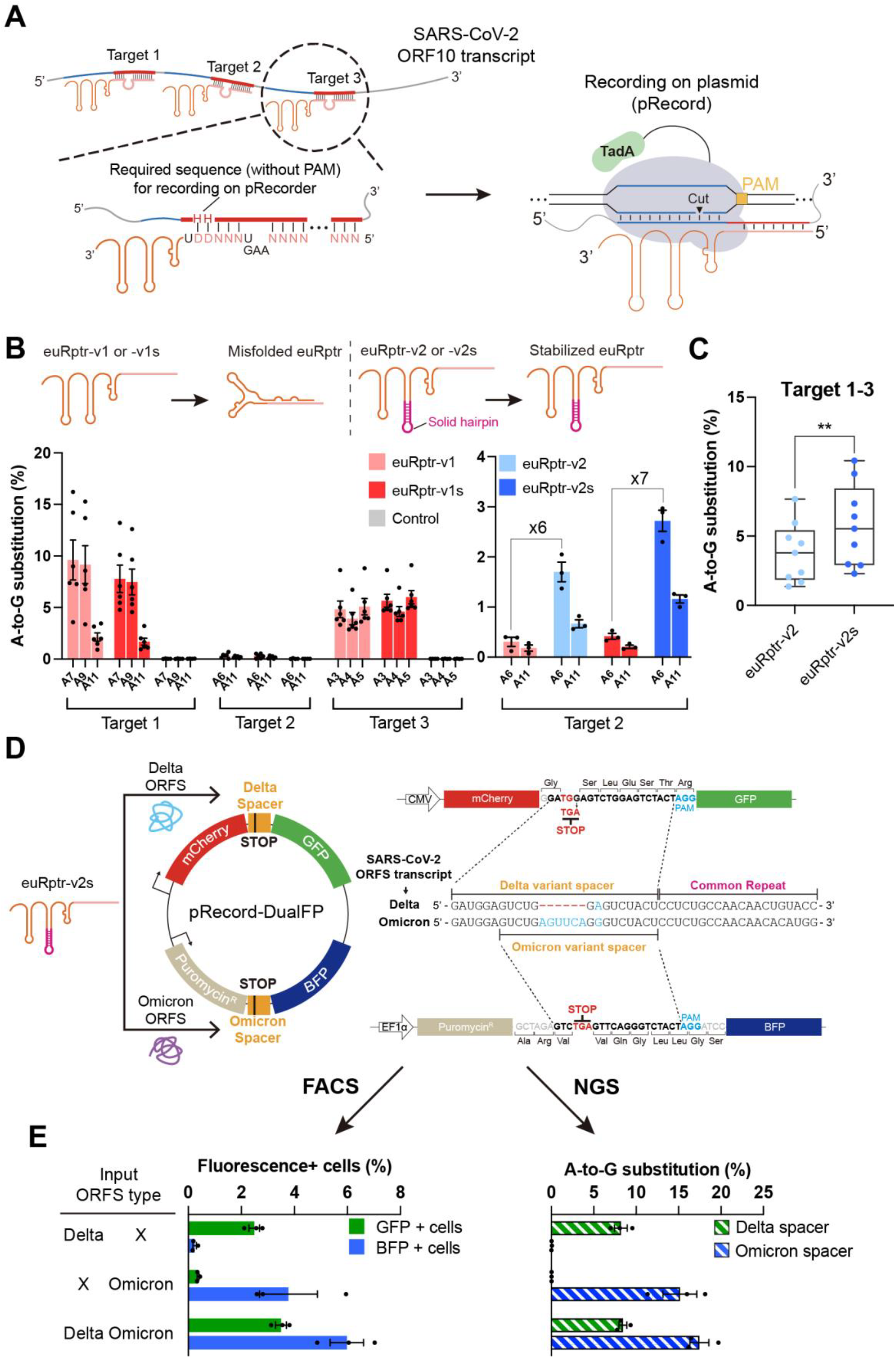
SARS-CoV-2 variant-specific RNA recording with CHEETAH and adenine base editors. **(A)** Schematic of the SARS-CoV-2 ORF10 transcript recording on pRecord. A total of three hijacking targets were designed using the 5’-NHHNNN-3’ rule. For RNA recording, NG-ABE8e was used, and the pRecord containing 5’-NG-3’ PAM sequence was prepared. **(B)** Recording frequencies across the three hijacking targets (*n* = 3; error bars represent mean ± SEM). The ORF10 transcript was overexpressed from the plasmid. The difference between euRptr-v1 and euRptr-v2 is described schematically. The fold change was calculated by comparing the average A-to-G substitution rate at A6 of target 2. **(C)** Comparison of euRptr-v2 and euRptr-v2s in targets 1-3 of ORF10. **, *P* = 0.0020 (paired Student’s *t*-test). **(D)** Schematic of SARS-CoV-2 variant-specific RNA recording with CHEETAH and NG-ABE8e. ORF transcripts of the Delta and Omicron (BA.2) variants were overexpressed from plasmids. Both variants share the same euRptr-v2s, which targets the common repeat region of the ORF transcripts. In the reporter pRecord, the corresponding target spacer containing a stop codon was embedded between two marker proteins. The mCherry protein serves as a positive control for transfection and the puromycin resistance gene prevents the alternative translation frame of BFP. Releasing the variant-specific stop codon results in corresponding fluorescence signal (Delta; GFP, Omicron; BFP). **(E)** The recording results were measured by fluorescence activated cell sorting (FACS) (left) and targeted deep sequencing (right) (*n* = 3; error bars represent mean ± SEM).

Next, we investigated the target discrimination ability of CHEETAH with euRptr-v2s for RNAs with high similarity. We evaluated two components that contribute to target discrimination: the repeat region of the euRptr and the spacer region of the target DNA. By performing mutagenesis of the lower stem and upper stem regions of the repeat region of euRptr-v2s targeting the ORF10 transcript fragment (**Fig. S8A**), we determined that more than two mismatches are required in the lower stem loop region for transcript-specific hijacking (**Fig. S8B**). The mutagenesis of the spacer region of euRptr-v2s (**Fig. S8C**) revealed that mismatched sequences should be positioned in the PAM-proximal seed region (1th-12th) for transcript-specific hijacking (**Fig. S8D**), which is similar to on-target discrimination requirements of the typical gRNA of SpCas9 (*38*).

The spike domain (ORFS) of SARS-CoV-2 Delta and Omicron variants (BA.2) have high sequence similarities, making it challenging to distinguish them using available diagnostic methods (*39, 40*). We predicted that CHEETAH could hijack both viral RNAs of Delta and Omicron variants with a shared-euRptr targeting the common repeat sequence of Delta and Omicron and specifically record their presence on Delta- or Omicron-specific pRecord plasmids (pRecord-ORFS). We designed ten euRptr-v2s candidates within the mutation-containing spike domain of the SARS-CoV-2 ORFS. Each euRptr targets a common repeat region and sequences specific to the Delta or Omicron variants are located upstream of each euRptr (**Fig. S9A**). Targeted deep sequencing of pRecord-ORFS revealed that five out of ten targets exhibited moderate base editing efficiencies (more than 1%) with NG-ABE8e for both RNA variants (**Fig. S9B**). We selected euRptr-v2s-9 for CHEETAH-mediated RNA recording. According to the euRptr-v2s-9 sequences, we prepared a specific pRecord (pRecord-DualFP, indicating a dual-fluorescence protein system), in which individual target sequences of both Delta and Omicron variants contain a stop codon (5’-TGA-3’) and the Delta sequence was located upstream of green fluorescent protein (GFP) and the Omicron sequence was upstream of blue fluorescent protein (BFP) (**Fig. 3D**). If the exogenous Delta or Omicron ORFS transcripts were detected in cells, euRptr-v2s-9 would hijack the transcripts and induce base-editing with NG-ABE8e on the pRecord-DualFP, modifying the corresponding stop codon and producing either GFP or BFP (**Fig. 3D**). Specifically, GFP would be produced when Delta transcripts were hijacked and recorded, whereas BFP would be produced when Omicron transcripts were hijacked and recorded. We transfected NG-ABE8e, euRptr-v2s-9, pRecord-DualFP along with Delta and Omicron ORFS expression plasmids into HEK293T cells. Three days after transfection, we detected green fluorescence from GFP in cells transfected with the Delta ORFS-overexpression plasmid and blue fluorescence from BFP in cells with the Omicron ORFS-overexpression plasmid (**Fig. S10**). Base editing occurred with a frequency of ∼7% at the Delta recording site and ∼15% at the Omicron recording site, when the matching variant was transfected into HEK293T cells (**Fig. 3E, right**). Consistent with the difference in base-editing frequencies, cells expressing the Omicron ORFS also tended to have a higher frequency of BFP positive cells quantified by fluorescence-activated cell sorting (FACS) compared to cells expressing the Delta ORFS (**Fig. 3E, left**). The signal for each output —base-editing and fluorescence frequency— matched the presence of the variant ORFS plasmid, indicating that CHEETAH selectively and specifically recorded the presence of each variant alone or in combination on a specifically designed pRecord-DualFP.

### CHEETAH records RNA-corresponding barcodes using prime editors

Base editing for recording is useful, but its application to genetic circuit design is limited due to its restricted editing scope, which primarily supports C-to-T or A-to-G substitutions (*11*). In contrast, prime editing is a more versatile tool capable of writing genetic information, including insertions, deletions, and all 12 types of substitutions (*23*). We evaluated the compatibility of CHEETAH with prime editing by developing a newly engineered version of euRptr, termed peuRptr, capable of prime editing after hijacking intracellular RNAs (**Fig. 4A**). Based on the euRptr-v2s, the peuRptr-v2s architecture includes a 3’ extension like a prime-editing gRNA and a pseudoknot (tevoPreQ1) at the 3’ end for preventing RNA degradation by 3’-5’ exonucleases in cells (*41*). To validate the activity of peuRptr-v2s, we tested two cases: one inserted an AT barcode and the other a CA barcode, both of which are designed to hijack the target 3 region of SARS-CoV-2 ORF10 transcript as shown in **Fig. 3A**. We transfected plasmids encoding the NG-PE2 with or without nicking gRNAs, each peuRptr-v2s, and ORF10 RNA transcripts, as well as pRecord-ORF10 into HEK293T cells. NG-PE2 is a variant of PE2 that recognizes NG-PAM (*23, 29*). Three days after transfection, targeted deep sequencing of pRecord revealed prime-editing efficiencies averaging 1.6% with PE2 and 2.6% with PE3 (i.e., addition of nicking gRNA) (**Fig. 4B**). These results suggested that CHEETAH can hijack exogenous RNAs using peuRptr-v2s, enabling the marking of corresponding barcode sequences on specific pRecord-ORF10 by PEs.

**Figure 4.**
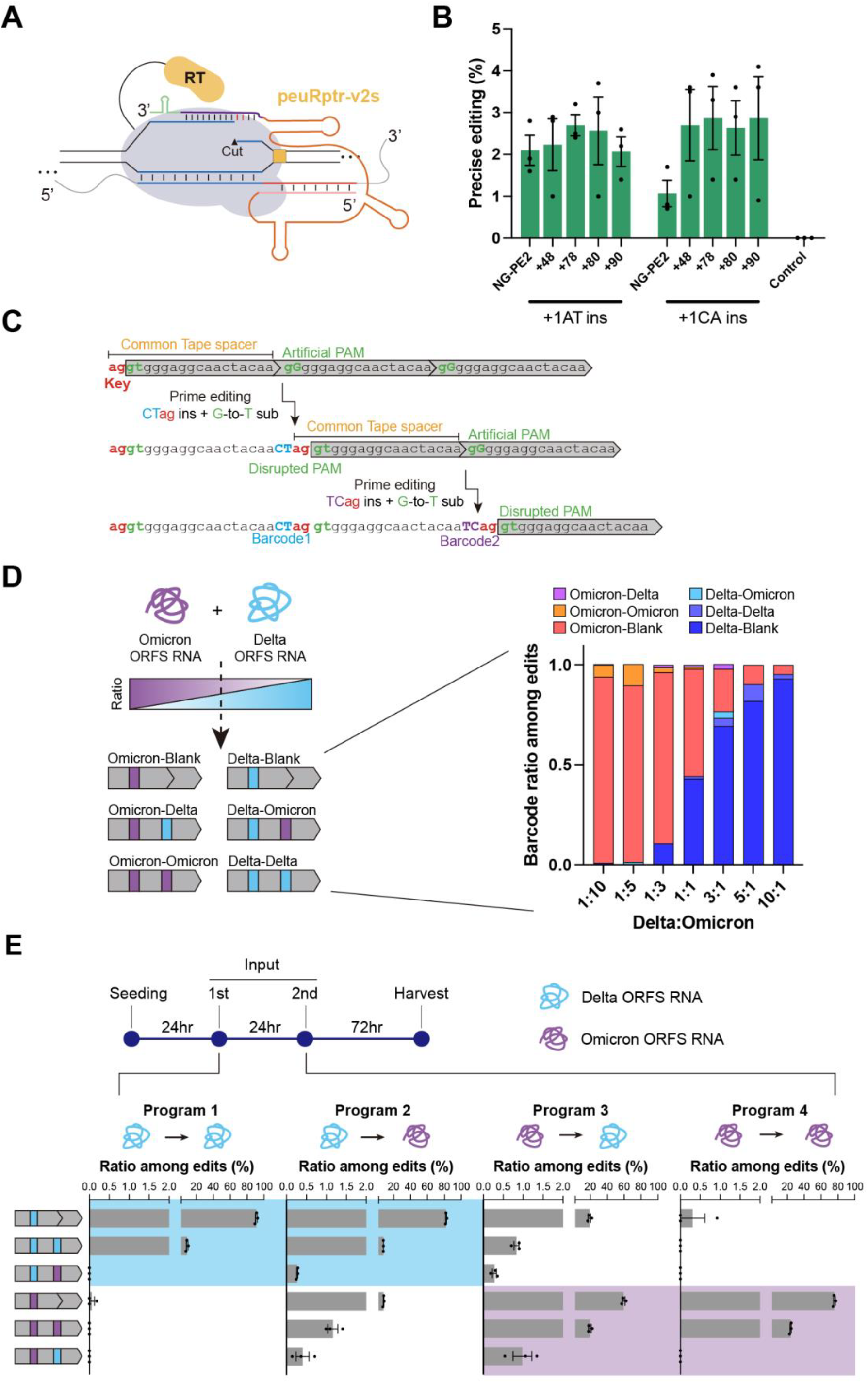
Viral RNA recording with CHEETAH and prime editors. **(A)** Schematic of CHEETAH using a prime editor. The peuRptr-v2s was designed by adding a 3’ extension and pseudoknot to euRptr-v2s. Data for NG-PE2 or NG-PE7 are shown in 4B or 4D-E, respectively. **(B)** ORF10 transcript recording frequencies on pRecord-ORF10. NG-PE2 was used with or without an additional nicking gRNA (*n* = 3; error bars represent mean ± SEM). Additional nicking position was described as a distance between peuRptr-induced nick and sgRNA-induced nick. **(C)** Schematic of two successive SARS-CoV-2 variant-specific RNA recording events on DNA Tape 2. The DNA Tape consists of a tandem repeat that has a 5’end truncated sequence (grey boxes). The 4 bp insertion (2 bp for barcode and 2 bp for key sequence) and 1 bp substitution (for disrupting the artificial NG PAM sequence) by prime editing activates the next repeat. Both transcripts of SARS-CoV-2 variants share the same DNA Tape but have variant-specific peuRptr-v2s inserting a variant-specific 2 bp barcode. **(D)** Multiplexed recording of ORF transcripts from two SARS-CoV-2 variants (Delta and Omicron) on DNA Tape 2. Different ratios of Delta and Omicron ORF expression plasmids with the corresponding peuRptr-v2s and NG-PE7 expression plasmids were transfected into the lentiviral DNA Tape-integrated HEK293T cell line. The ratio of barcodes inserted into DNA-Tape were determined from both unigram (variant-blank) and bigram (variant-variant) insertion events (*n* = 3). **(E)** Time-resolved recording of Delta and Omicron ORFS transcripts. Plasmids expressing NG-PE7, peuRptr-v2s, and ORF transcripts were transfected for each of the 2 inputs; the pRecord-Tape was provided only during the first input transfection. The ratio of each DNA Tape pattern was calculated among the entire edits (*n* = 3; error bars represent mean ± SEM). The blue box and purple box indicate DNA Tape patterns of the Delta and Omicron barcode at position 1, respectively. The blue box is symmetric with the purple box due to the difference in the first input.

Given the compatibility of CHEETAH with prime editing, we examined whether it could function as a DNA-Typewriter, a strategy enabling multiplexed and time-resolved recording in human cells (*42*). Unlike the recording strategy using base editors in **Fig. 3D**, we hypothesized that CHEETAH could hijack either Delta or Omicron ORF variants through variant-specific peuRptrs and distinctively record variant-specific barcodes on a common pRecord-Tape using PEs. We designed two DNA Tapes (DNA Tape 1 and 2) using two identical sequences between Delta and Omicron ORF transcripts (**Fig. S11A** and **S11B**). The upstream identical sequences served as a common DNA Tape spacer, and downstream distinct sequences served as variant-specific repeat regions for hijacking by peuRptrs. The DNA Tapes consist of three components: (i) the “key” sequence of a 2∼4 bps at the 5’ end of the common Tape spacer, which should be included as a desired prime edit along with a transcript-specific barcode; (ii) a tandem array of repeated units serving as imperfect spacers; (iii) an artificial PAM that is a modified sequence within the tandem array that functions as a PAM for NG-PE. Briefly, inserting the key sequence and introducing the artificial PAM results in an intact common Tape spacer, enabling subsequent rounds of prime editing (**Fig. 4C** and **S11B**).

Using two different dedicated DNA Tapes (DNA Tape 1 and 2), we tested unigram recording efficacy by co-transfecting plasmids expressing NG-PE7 (*43*), variant-specific peuRptr-v2s, and ORF transcripts. Depending on the input source, the corresponding barcode was correctly inserted into all types of DNA Tape, with average prime editing rates of three replicates as follows: 1.35% for Delta barcode and 1.71% for Omicron barcode in DNA Tape 1, 0.86% for Delta BC and 1.58% for Omicron BC in DNA Tape 2 (**Fig. S11C**). To validate CHEETAH’s functionality as a DNA-Typewriter for multiplexed and time-resolved recording, we selected DNA Tape 2 (**Fig. 4C**), which showed moderate insertion frequency with two-nucleotide barcodes. For multiplexed recording, we recorded mixtures of SARS-CoV-2 ORF transcripts with varying Delta:Omicron input ratios (1:10, 1:5, 1:3, 3:1, 5:1 or 10:1) in HEK293T cells bearing stable knock-in of DNA Tape 2 target arrays. Although overall editing rates were modest (only with average insertion frequencies of 2.58% for a 1:10 ratio, 1.61% for 1:5, 1.01% for 1:3, 1.60% for 1:1, 1.46% for 3:1, 1.74% for 5:1, and 1.19% for 10:1), as expected, the barcode frequency on the integrated DNA Tape 2 reflected the input ratio (**Fig. 4D**).

For time-resolved recording, we designed four different event programs, i.e., program 1, 2, 3, and 4 indicating Delta-Delta, Delta-Omicron, Omicron-Delta, and Omicron-Omicron, respectively. All possible orders were considered for two sequential inputs to pRecord-Tape (DNA Tape 2), provided 24 hours apart, with cells harvested 72 hours after the last input (**Fig. 4E**). We confirmed that the sequencing results reflect the correct sequential recording program. Because the first input source was recorded at position 1 of the DNA Tape with a higher frequency, the ratio of edited DNA Tape patterns in Programs 1 and 2 are symmetrically reflected in Programs 4 and 3, respectively. We could also distinguish between homogeneous and heterogeneous programs through the information of both barcode types in position 1 and the ratios of heterogeneous patterns in DNA Tapes (**Fig. 4E**).

Taken together, we validated that CHEETAH is compatible with PE through the use of a peuRptr-v2s. Furthermore, we showed that CHEETAH enables multiplexed and time-resolved recording of RNAs on a DNA Tape, functioning as a DNA-Typewriter system.

## Discussion

In this study, we demonstrated that euRptr can hijack cellular RNAs to recruit CRISPR-associated complexes to genomic or plasmid DNA in human cells. By characterizing the mutation tolerance of both SpCas9 and NG-SpCas9, we determined that NG-SpCas9 exhibits greater tolerance for modifications in the secondary RNA structure between the crRNA repeat and tracrRNA anti-repeat regions than the canonical SpCas9. Based on this information, we extracted a sequence formula for euRptr for application of NG-SpCas9 or NG-ABE8e in human cells. To induce gene editing or base editing on the corresponding endogenous target sites with NG-SpCas9 or NG-ABE8e, respectively, target sites of euRptr must contain 5’-NGHNNNNN-3’ sequences that functions as an NG PAM on the gene. Then, the target sites within mRNA from this gene interacts with the euRptr. For the abundantly expressed genes in HEK293 cells, we observed low but certain editing outcomes, indicating a successful euRptr-leveraged self-editing.

To record RNA expression events on a specific pRecord of DNA, a different formula for euRptr was necessary. Target sites must contain 5’-NHHNNNNN-3’ sequences lacking an NG PAM to avoid self-editing by the euRptr. Through this strategy, we demonstrated that a shared-euRptr can recognize different RNA fragments of SARS-CoV-2 Delta or Omicron variants and recruit ABEs to record the presence of the RNAs on a specifically designed pRecord. By engineering peuRptr, we further proved that CHEETAH is compatible with PEs in human cells, thereby expanding the capability of the system. By adopting the DNA-Typewriter approach, we demonstrated that CHEETAH can induce two-base barcode insertions on DNA tape enabling multiplexed and time-resolved recording of different SARS-CoV-2 Delta or Omicron variants. Thus, CHEETAH is a promoter-independent system for detecting transcripts. These strategies that use base editing or prime editing can be utilized for record intracellular RNA expression events during cell division, cell differentiation, viral infection, or lineage switching in eukaryotic cells in the future.

Several methods monitor mammalian cell status through tracking gene expression, however each has limitations. Approaches based on adenosine deaminase acting on RNA (ADAR) enzymes detect a specific target RNA but the detected signal is transient and is not permanently recorded in cells (*44-46*). DNA-based recording strategies enable the creation of a permanent record on DNA, as a memory device, but they typically rely on promoters responsive to specific types of signals, limiting the number and diversity of input signals (*9-13, 42*). In addition, previous promoter-independent RNA-recording strategies that use RT-Cas1-Cas2 or Rptr record specific RNA expression events on DNA but are limited to prokaryotes (*17-19*). Our study provides a versatile DNA-based RNA-recording system (CHEETAH) for monitoring RNA expression events in eukaryotic cells.

### Limitation of the study

Despite the high potential of CHEETAH in self-editing or RNA recording in human cells, further studies are required to improve the general recording efficacy and activity with intracellular RNAs by increasing the efficiencies of CRISPR-associated tools, utilizing other CRISPR-Cas effector with expanded PAM sequences, and engineering euRptr sequences to hybridize with target RNAs more strongly. Enhancing the CHEETAH-mediated RNA-recording activity, particularly for endogenously expressed mRNAs that are relatively low concentrations in cells, would be crucial for enabling biological applications, such as investigating disease-causing mRNA splicing variants and lineage tracing.

## Materials and Methods

### Plasmid construction

Ligation cloning was performed to obtain an acceptor plasmid of the euRptr expression vector and of the episomal target plasmid (pRecord). For euRptr expression cloning, pRG2 (Addgene no. 104174) plasmid was digested with BsaI (New England Biolabs, R3733L) and SpeI (New England Biolabs, R3133S). For pRecord cloning, pDAS12124_PEAR-GFP-preedited (Addgene no. 177179) plasmid was digested with PciI (New England Biolabs, R0655S) and MluI (New England Biolabs, R3198S). The required DNA oligonucleotides were ordered from Macrogen (Korea) and annealed to produce double-stranded DNA followed by 5’-phosphorylation using T4 PNK (Enzynomics, M005L). Digested vector fragments and 5’-phosphorylated dsDNA fragments were ligated using T4 DNA ligase (Enzynomics, M001L) (24 hrs at 25 °C).

Gibson assembly was performed to obtain the SARS-CoV-2 variant-specific fluorescence expression plasmid (Reporter plasmid) and the NG-PE7 expression plasmid. For reporter plasmid cloning, p3S-Cas9-HN (Addgene no. 104171) was digested with HindIII and XhoI. To prepare the insert DNA fragments, mCherry region and copGFP region were amplified from dCas9-VPR_P2A_mCherry (Addgene no. 154193) and pAAVS1P-iCAG.copGFP (Addgene no. 66577). The puromycin resistance gene and TagBFP regions were amplified from pAX198 (Addgene no. 173042). For NG-PE7 expression plasmid cloning, pCMV-PE7 (Addgene no. 214812) plasmid was digested with SacI (New England Biolabs, R3156S) and BamHI New England Biolabs, R3136S). To prepare the insert DNA fragments, the NG-Cas9 region and RT region were amplified from SpCas9-NG plasmid (Addgene no. 138566) and pCMV-PEmax (Addgene no. 174820). All insert DNA fragments were amplified with Q5 hotstart High-Fidelity DNA polymerase (New England Biolabs, M0493L). Digested vector fragments and insert DNA fragments were mixed in a volume of 10 μl containing 2U T5 exonuclease (New England Biolabs, M0363S), 12.5U Phusion DNA Polymerase (Thermo Fisher Scientific, F530L), 2kU Taq DNA ligase (New England Biolabs, M0208S), 0.2 M Tris-HCl (pH 7.5), 0.2 M MgCl_2_, 2 mM dNTPs, 0.2 M dithiothreitol, 25% PEG-8000 and 1 mM NAD, and were incubated at 50 °C for 6 hrs.

### RT-qPCR analysis

NucleoSpin RNA Plus Kits (MACHEREY-NAGEL, 740984.250) was used for total RNA extraction following the manufacturer’s protocol. cDNA was synthesized by adding 1 μg of RNA sample with the reagents of ReverTra Ace-α-(TOYOBO, FSK-101F) and incubated for 30 °C 20 min, 42 °C 60 min, 99 °C 5 min. Synthesized cDNA (2 μl) was used for qPCR template. iTaq Universal SYBR Green Supermix (Bio-rad, BR1725121) and qPCR instrument (Bio-rad, CFX96) were used for this experiment.

### Transfection of mammalian cells

HEK293 (ATCC CRL-1573) and HEK293T (ATCC CRL-3216) cells were cultivated in DMEM supplemented with 10% fetal bovine serum (FBS) and 1% ampicillin in an incubator at 37 °C with 5% CO_2_. Before transfection, 0.5 x 10^5^ cells per each well were seeded in 48-well plates. For endogenous target editing, Cas nuclease, ABE, or PE expression plasmids (375 ng) and sgRNA or euRptr expression plasmids (125 ng) were mixed with 0.5 μl jetOPTIMUS reagent (Polyplus, 101000006). For experiments with additional expression of RNA fragments, 125 ng of RNA fragment expression plasmids were additionally added and 0.6 μl jetOPTIMUS reagent were used. For episomal target recording, the pRecord plasmids (150 ng) were additionally added with other plasmids expressing ABE or PE, euRptr, and RNA-of-interest, and 0.75 μl jetOPTIMUS reagent were used. The media was changed with fresh DMEM 24 hr after transfection. Genomic DNA was isolated 5 days after transfection.

### Lentiviral infections for knock-in cell line generation

Lentivirus was produced in the Lenti-X 23T cell line (Takara Bio). Transfection was carried out with psPAX2 (Addgene #12260), pMD2.G Addgene #12259), and NG-PE7 expression plasmids using Lipofectamin 2000 (Invitrogen, 11668019). The cultured medium was replaced one day post-transfection. Lentivirus-containing medium was harvested 48 hrs and 72 hrs after transfection and filtered through a 0.45-µm syringe filter. The lentivirus was concentrated using the PEG-it virus precipitation solution (System Bioscience, LV825A-1). The viral titer was determined by performing lentiviral transductions at varying concentrations in a 48-well plate format. After titration, the lentiviral library was aliquoted and stored at -80 °C.

### Flow cytometry

Transfected cells were collected through centrifugation (500 g, 3 min at 4 °C). After discarding the supernatant, the cell pellet was resuspended in flow cytometer buffer (DPBS, 2% FBS) followed by centrifugation (500 g, 5 min at 4 °C). After discarding the supernatant, the cell pellet was resuspended in flow cytometer buffer again and transferred into the flow cytometer tube with a cell-strainer cap (FALCON, 352235). Fluorescence was measured by BD FACS Canto II (Becton, Dickinson and Company) instrument and analyzed by FlowJo v.10 software.

Cell sorting was performed using Cell Sorter (SONY, SH800S). For cell sorting, 1 x 10^7^ cells were collected using the same procedure described for flow cytometry cell preparation. Cells exhibiting the top 10% of fluorescence intensity were sorted in bulk and used for further experiments.

### Confocal imaging

HEK293T cells were seeded in confocal dishes (SPL, 214350) and transfected with total 1 μg plasmid DNA. Imaging was performed 48 hrs after transfection with the LSM980 (ZEISS) with Airyscan2. The excitation laser wavelengths were 405 nm (TagBFP), 488 nm (copGFP), and 561 nm (mCherry). All images were acquired using a 20x magnification, and the focus was adjusted based on the mCherry protein. Image processing was performed using ZEISS ZEN 3.9.

### Targeted deep sequencing

Cells were pelleted by centrifugation. The cell pellet was resuspended in 100 μl proteinase K extraction buffer [40 mM Tris-HCL (pH8.0), 1% Tween-20, 0.2 mM EDTA, 10 mg proteinase K, 0.2% Nonidet P-40 (VWR, 97064-730)] and incubated at 60 °C for 15 min followed by an incubation at 98 °C for 5 min. Genomic DNA (2 μl) was used as a PCR template for PCR amplification (98°C 10 sec, 56 °C 10 sec, and 68 °C 30 sec for 30 cycles) using KOD-Multi&Epi (TOYOBO, KME-101). The resulting amplicon (1 μl) was used for further PCR amplification with the same PCR conditions and a next generation sequencing library was prepared. The sequencing library was analyzed using an Illumina Miniseq instrument.

### Analysis of editing outcome

Indel, base editing, and prime editing frequencies were analyzed using Cas-Analyzer (http://www.rgenome.net/cas-analyzer/) (*47*), BE-Analyzer (http://www.rgenome.net/be-analyzer/) (*48*), and PE-Analyzer (http://www.rgenome.net/pe-analyzer/) (*49*), respectively. The analysis code for recording on DNA Tape is available at (https://github.com/BaeLab/CHEETAH/tree/main).

## Data availability

Targeted deep sequencing data have been deposited in the NCBI Sequence Read Archive database (https://www.ncbi.nlm.nih.gov/sra) under accession number PRJNA1178377.

## Acknowledgements

This study was inspired by previous research conducted at the Chase L. Beisel lab. The authors thank Nancy R. Gough (BioSerendipity, LLC) for English-language editing. Most analysis of sequencing data was carried out using the computing server at the Genomic Medicine Institute Research Service Center. This work was supported by the Samsung Research Funding & Incubation Center of Samsung Electronics under Project Number SRFC-MA2101-06 to S.B.

## Author contributions

S.-H.L. and S.B. conceived this project; C.J. developed bioinformatics algorithms; S.-H.L. and

J.P. performed cell experiments; S.B. supervised this project; S.-H.L. and S.B. wrote the manuscript with the help of all other authors.

## Additional information

Supplementary Information accompanying this paper is available at http://

## Declaration of interests

S.-H.L., J.P., and S.B. have filed a patent application based on this work. The remaining authors declare no competing interests.

## Notes

### Competing Interest Statement

The authors have declared no competing interest.

