## Supplementary Material for "Hijacking and recording intracellular RNAs in human cells using eukaryotic reprogrammed tracrRNAs"

### Supplementary Materials

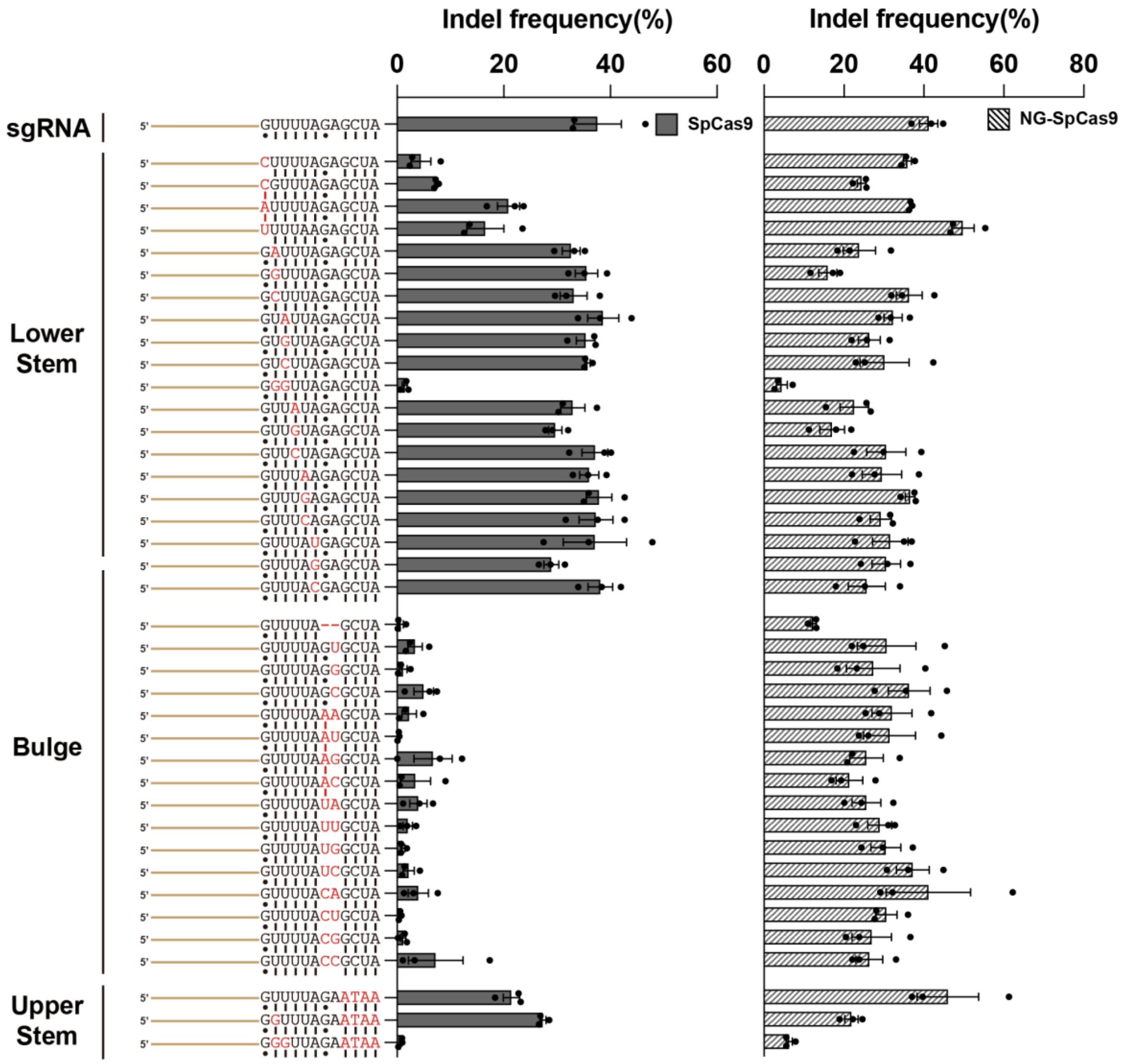


**Fig. S1. Indel frequencies generated by mutated sgRNA scaffold along with SpCas9 and NG-SpCas9.** The histograms show the raw data used in Figure 1B-C (*n* = 3; error bars represent mean ± SEM). Substitutions or deletions were applied to the lower stem, bulge, and upper stem of *TBP* gene-targeting sgRNA. For SpCas9, the positions of RNA secondary structure are vulnerable to the mutation, whereas NG-SpCas9 showed higher tolerance in same positions. Only 5’-NGG-3’ mutation in the lower stem or deletion of bulge (5’-GA-3’) disrupted the indel frequency of NG-SpCas9. Mutations are indicated in red. The upper stem mutations (ATAA) correspond to the same positioned sequence in the *TBP* target site. For base pairing annotation, dots indicate wobble base pairs, and dash lines indicate Watson-Crick base pairs.

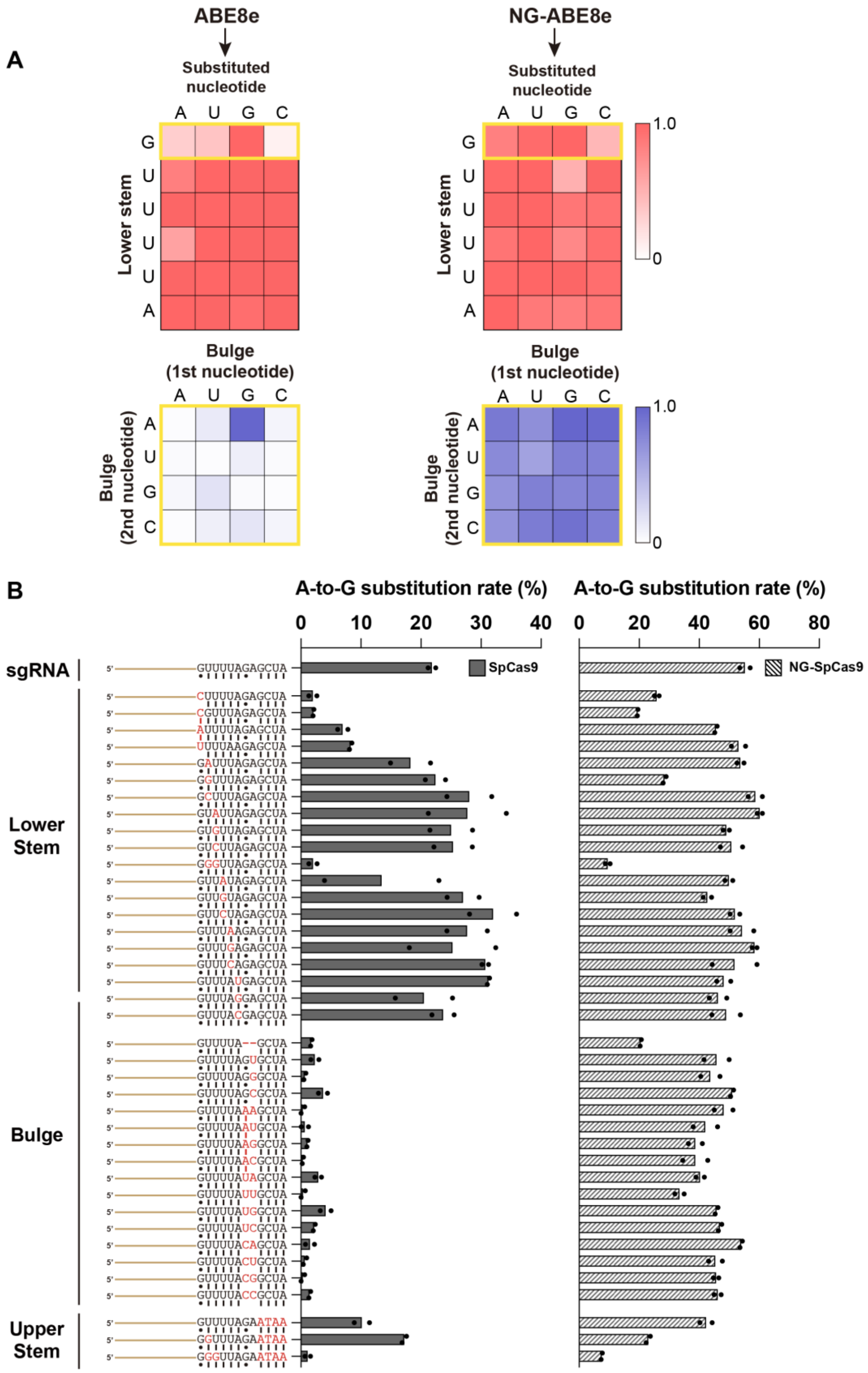


**Fig. S2. Evaluation of sgRNA scaffold tolerance for ABE8e and NG-ABE8e. (A)** The effect of simple substitutions on the lower stem and bulge regions of *TBP* gene-targeting sgRNA.

The A8-to-G substitution rate was measured in *TBP* target site. The nucleotide notation in the heatmap indicates the nucleotide in the repeat sequence. In the lower stem (5'-GUUUUA-3'), each nucleotide was independently substituted with all other nucleotides. In the bulge (5'-GA-3'), every combination of substitutions was tested. All heatmap values represent the fold change relative to wild-type sgRNA. Yellow boxes indicate positions related to RNA secondary structure. **(B)** The histograms show the raw data used to generate the heatmaps in A ($n$ = 2). For base pairing annotation, dots indicate Wobble base pairs, and dash lines indicate Watson-Crick base pairs.

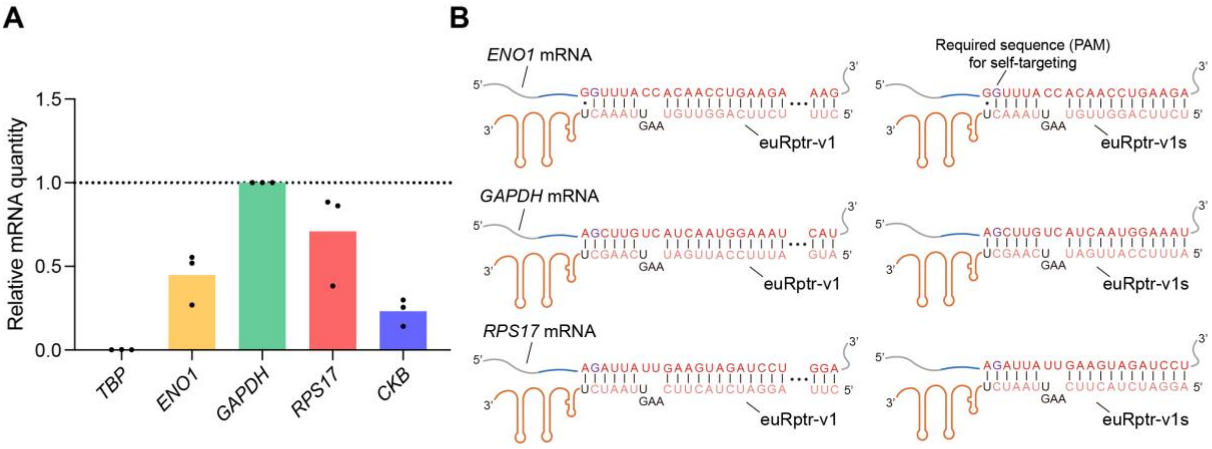


**Fig. S3. Investigation of endogenous mRNA abundance in HEK293 cell line. (A)** The total RNAs were extracted from three different dishes containing HEK293 cell lines. The total RNAs were reverse transcribed for cDNA generation. Then, qPCR was performed to measure the mRNA abundance. The quantified mRNA was normalized to *GAPDH* mRNA, with data shown for $n$ = 3 biologically independent samples. **(B)** Sequence information of endogenous mRNAs (*ENO1*, *GAPDH*, and *RPS17*) and euRptrs used in Figure 2B-2D and Figure S4 for euRptr-leveraged self-targeting. Target sites were selected to satisfy the 5’-NGHNNNNN-3’ rule. The purple-colored “G” on mRNA indicates the required sequence (PAM) for self-targeting.

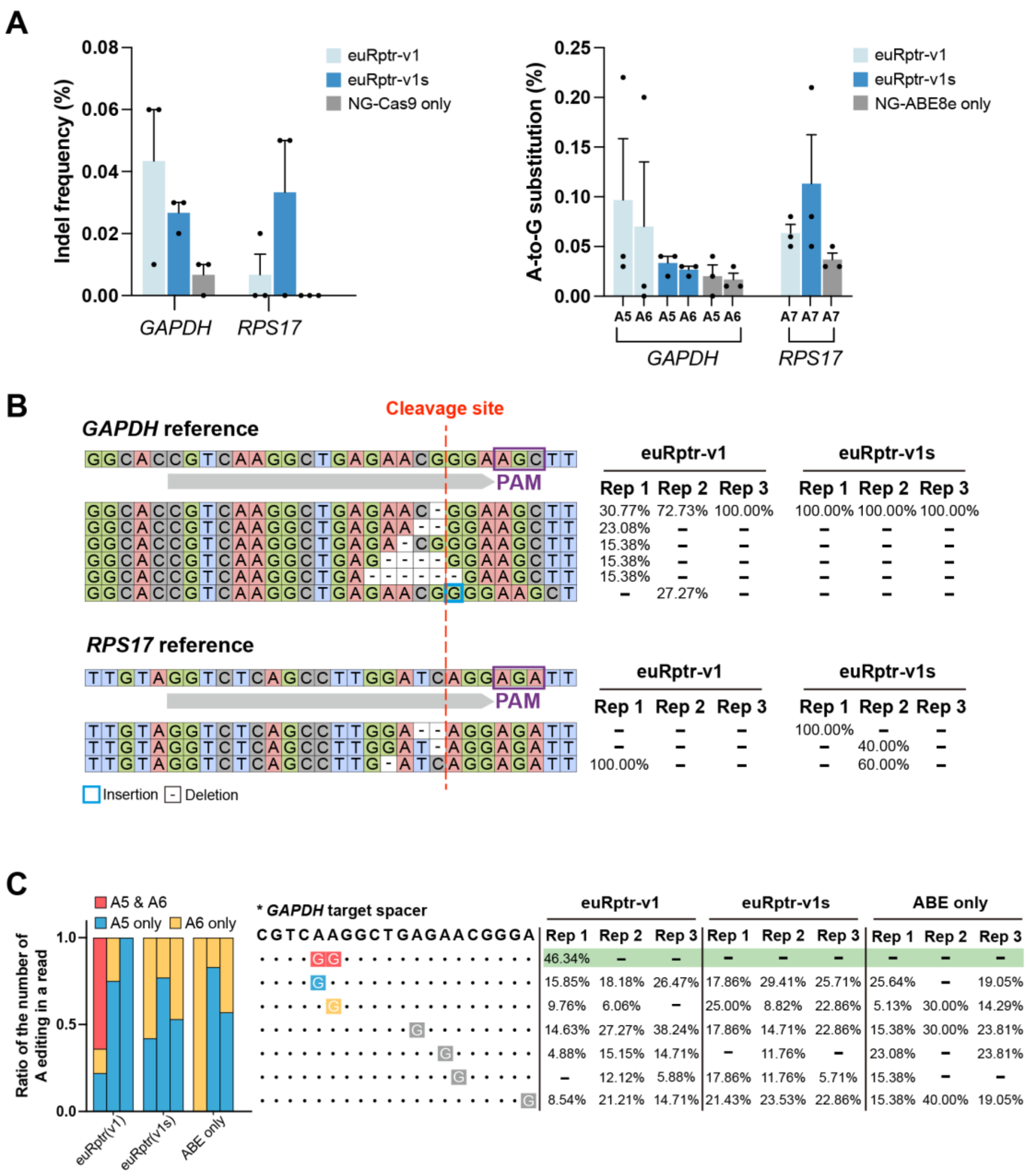


**Fig. S4. Genomic self-targeting via *GAPDH* and *RPS17* endogenous mRNAs hijacking with euRptr. (A)** Both euRptr and NG-SpCas9 or NG-ABE8e were overexpressed from plasmids. For the self-targeting, the rule (5'-NGHNNN-3' from downstream of the spacer) was applied to search for endogenous targets. The sequence should be in the exon, and the 5'-NGH-3' sequence acts as a PAM for the self-targeting. Indel frequency (left) and A-to-G substitution rate (right) at the *GAPDH* and *RPS17* endogenous target sites are shown in this figure ($n$ = 3; error bars represent mean ± SEM). **(B)** Indel patterns generated by NG-SpCas9 using hijacked endogenous *GAPDH* and *RPS17* mRNA. (**C**) Base-editing patterns generated by NG-ABE8e. In both B and C, the numbers indicate the percent ratio of each editing pattern among all mutations.

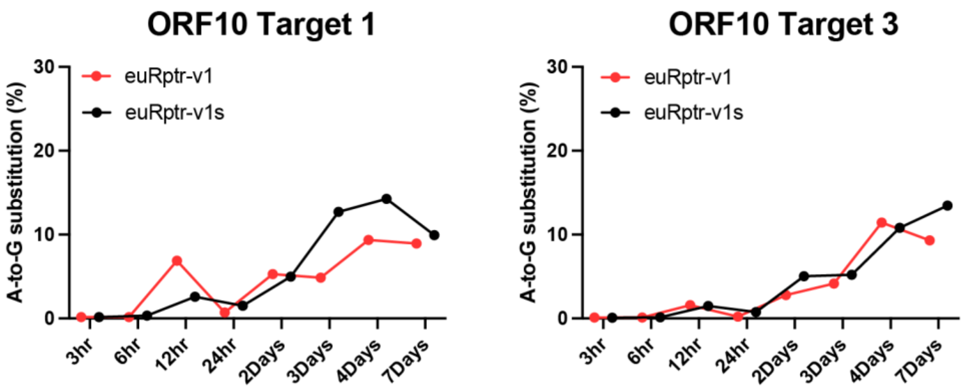


**Fig. S5. Recording profile on the episomal target plasmid over time.** ORF10 transcript recording on the pRecord-ORF10 using NG-ABE8e accumulated up to day 7. ORF10 transcript, euRptr, and NG-ABE8e expression plasmids were transfected in HEK293T cells. Data represent the mean value from *n* = 3 biologically independent experiments.

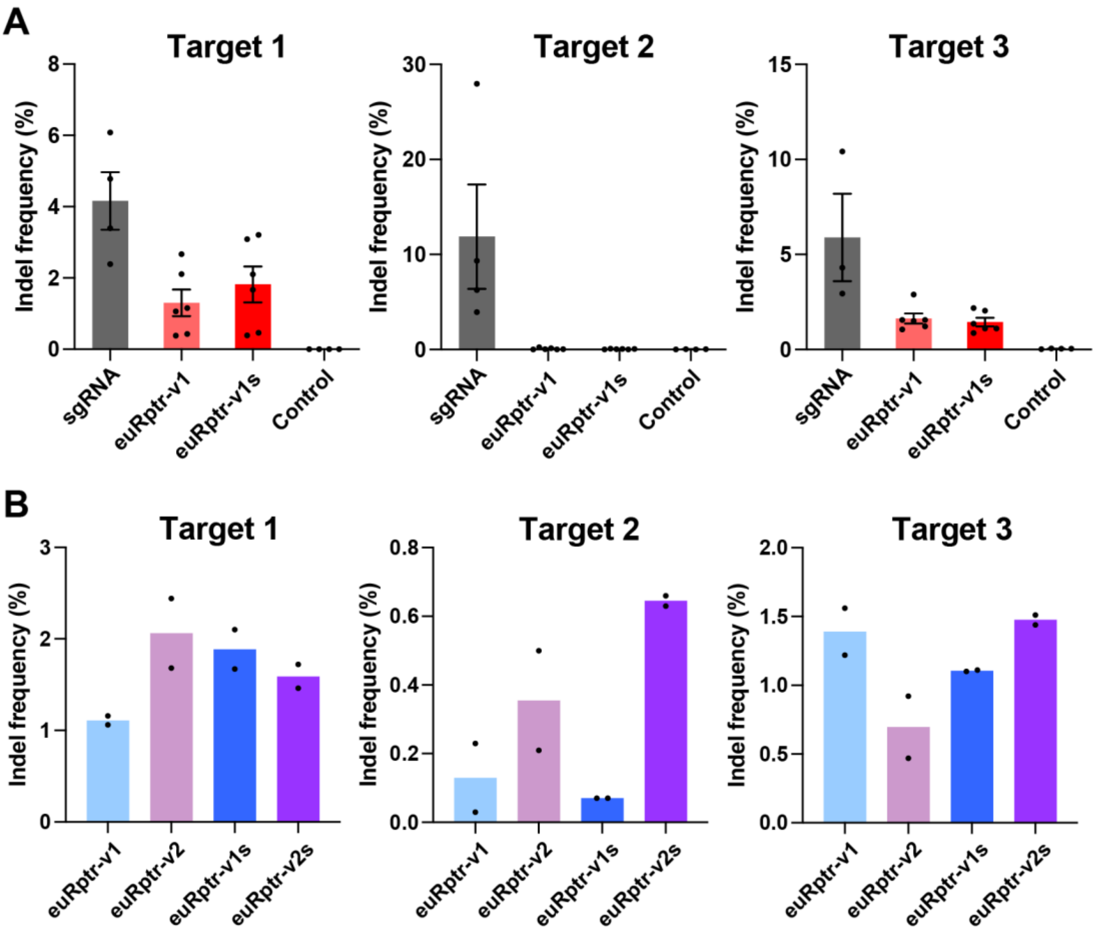


**Fig. S6. Target dependency of ORF10 transcript hijacking and editing by NG-SpCas9. (A)** ORF10 transcript, euRptr -v1 or -v1s, and NG-SpCas9 expression plasmids were transfected into HEK293T cells. Indel frequencies on pRecord-ORF10 were measured for ORF10 transcript recording for three target sites ($n > 3$; error bars represent mean ± SEM). **(B)** Conditions were the same as in A except euRptr secondary structure was stabilized with a solid hairpin in the hairpin region (euRptr-v2 and euRptr-v2s) ($n = 2$).

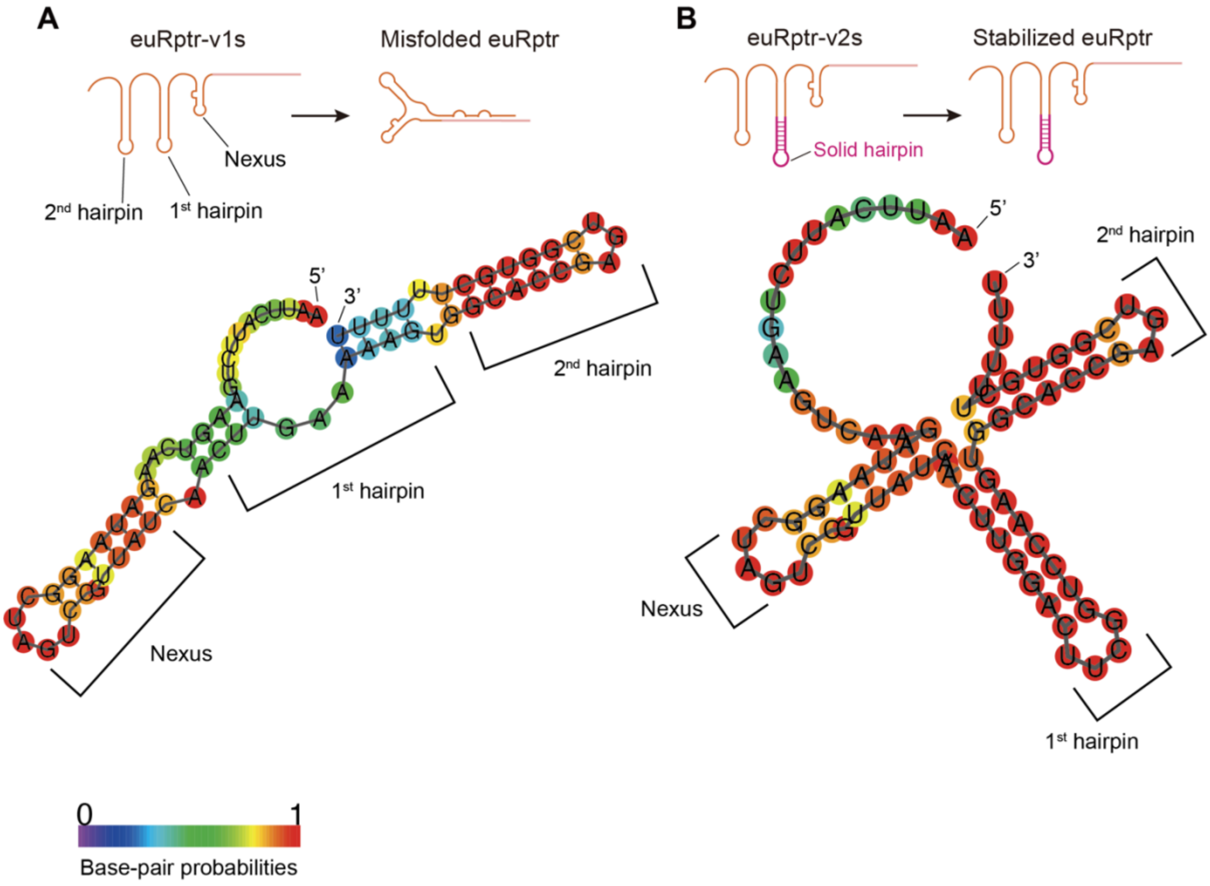


**Fig. S7. RNA secondary prediction of euRptr-v1s and euRptr-v2s targeting ORF10-target 2. (A-B)** Predicted RNA secondary structure of euRptr-v1s (A) and euRptr-v2s (B) targeting ORF10-target 2. The lower stability of the 1st hairpin of euRptr-v1s can result in misfolding that impairs hijacking of RNA-of-interest. A solid hairpin has high melting temperature and prevents misfolding. RNA secondary structure analysis was performed with RNAfold WebServer. The displayed structures indicate minimum free energy (MFE) structure and show the encoding base-pair probabilities.

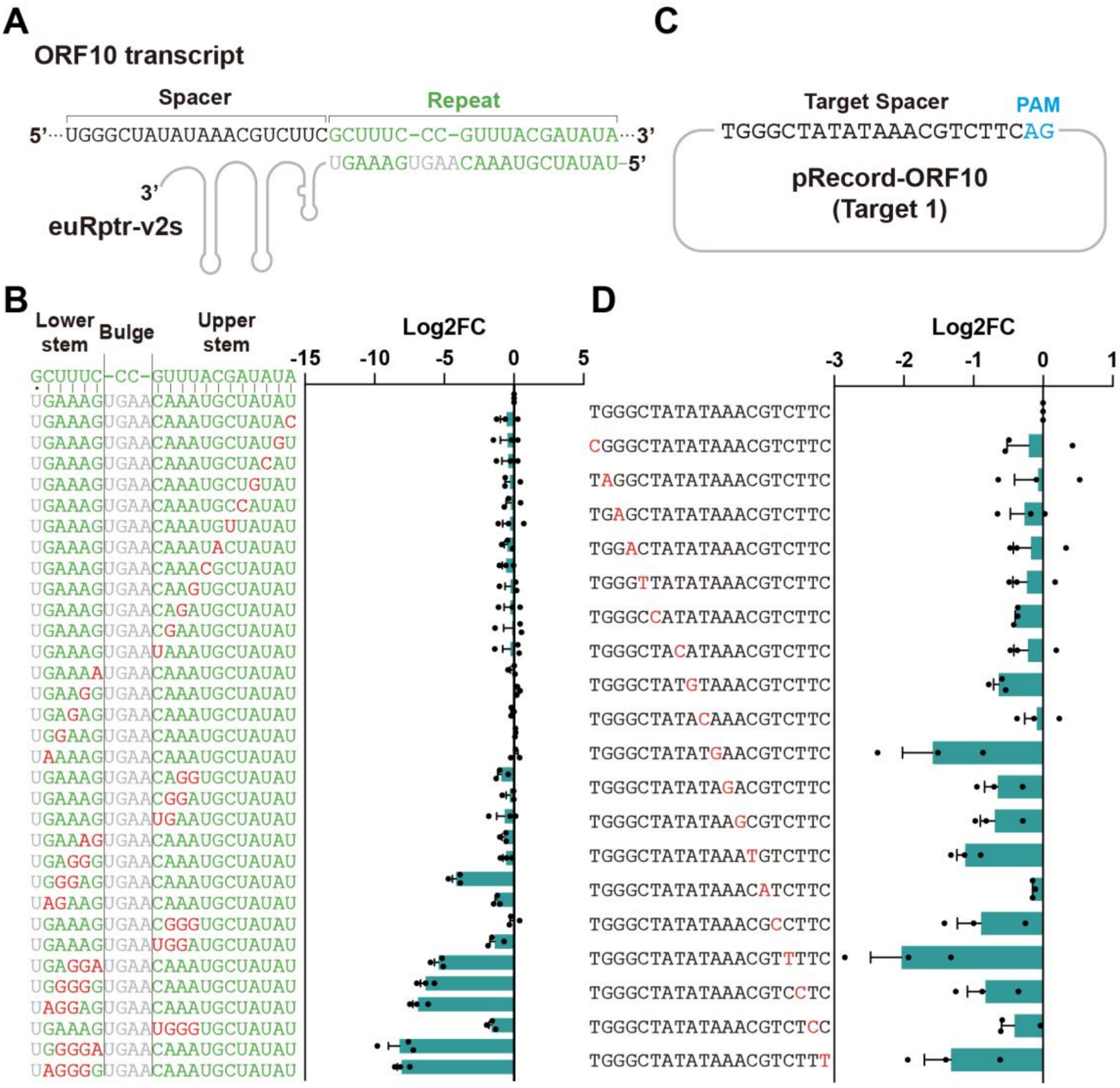


**Fig. S8. Investigation of discrimination ability of CHEETAH. (A)** Schematic representation of the repeat:anti-repeat region contributing to specificity. Similar non-target RNA could be discriminated through repeat:anti-repeat hybridization region. **(B)** Discrimination ability through anti-repeat region mismatches in euRptr-v2. Data were normalized to original euRptr-v2s ($n$ = 3; error bars represent mean ± SEM). **(C)** Schematic representation of the spacer region of pRecord which is the hybridization with the complement sequence of target RNA. Similar non-target RNA could be discriminated through spacer-DNA:hijacked-RNA hybridization region. **(D)** Evaluation of discrimination ability through mutations of spacer region. Mismatches in the PAM-proximal regions enable to distinguish the target RNA from the similar non-target RNA. Data was normalized to wild-type spacer sequence on episomal target plasmid ($n$=3; error bars represent mean ± SEM). Red letters indicate mismatch-inducing mutations.

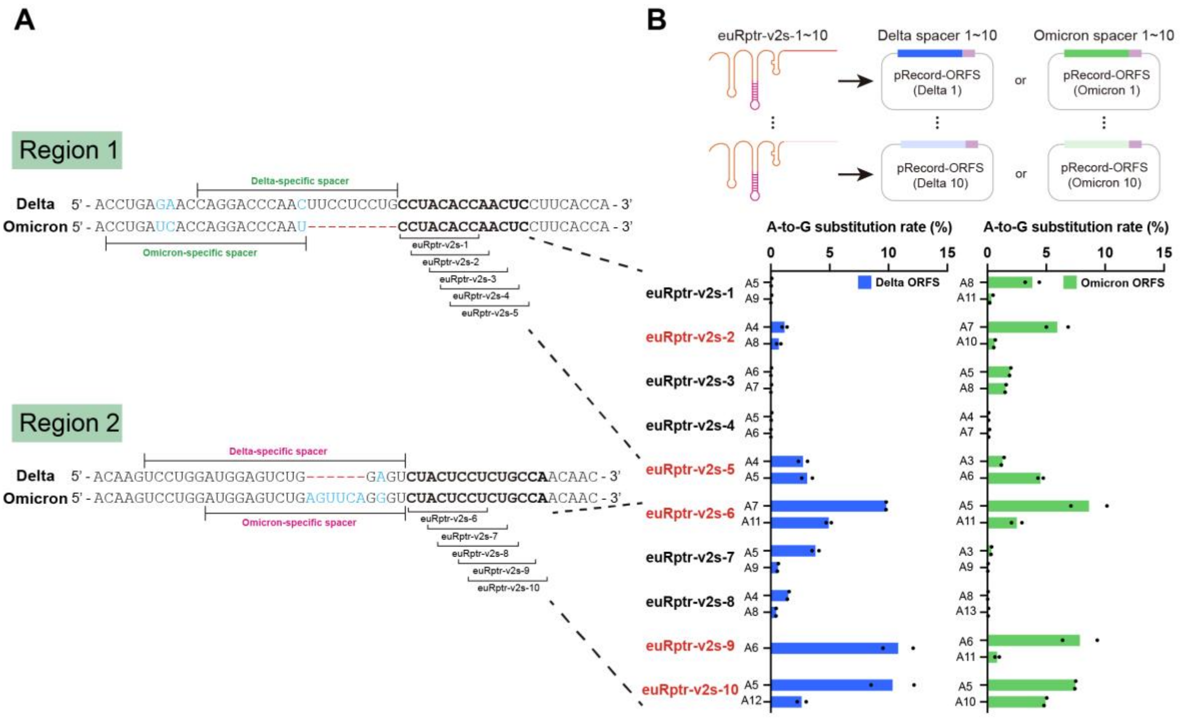


**Fig. S9. Evaluation of editing frequencies from similar targets for SARS-CoV-2 variant-specific recording with CHEETAH and adenine base editors. (A)** Two selected regions with sequence differences between Delta and Omicron ORF transcripts. Blue letters and dash lines indicate the substituted sequences and deleted sequences between two variants, respectively. Five targets for each region were designed for hijacking by euRptr-v2s. **(B)** Editing frequencies at all designed target sites (*n* = 2). The ORF transcripts, euRptr-v2s, and NG-ABE8e expression plasmids were transfected into HEK293T cells. The A-to-G substitution rate on pRecord-ORFS was measured to evaluate the recording frequency of each target site.

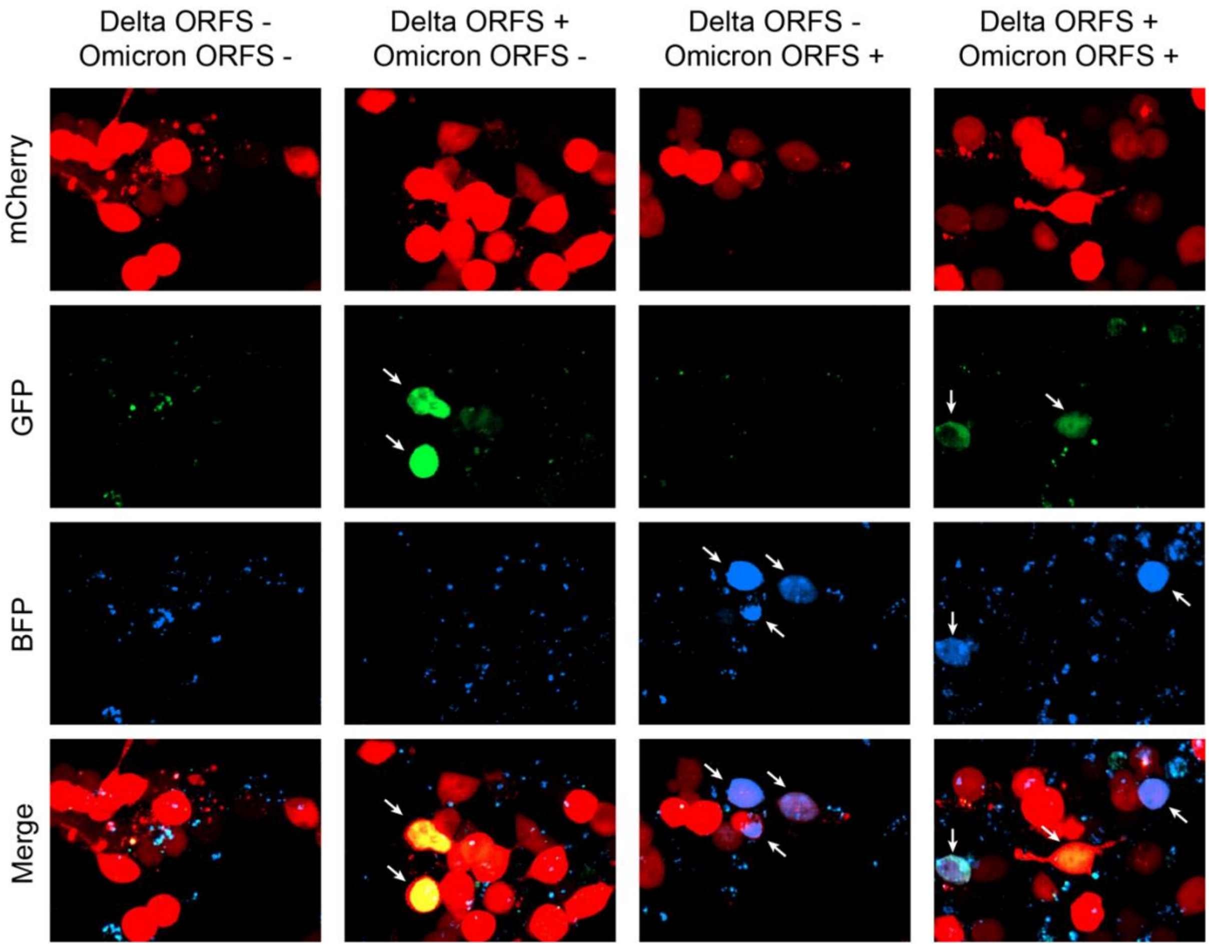


**Fig. S10. Confocal imaging results for variant-specific recording with CHEETAH and NG-ABE8e.** Imaging results of variant-specific recording using the same reporter pRecord in Figure 3D. Same experimental strategies with figure 3D-E was applied to this experiment. mCherry denotes the pRecord-reporter transfected cell, GFP and BFP denotes the presence of Delta and Omicron ORFS transcripts, respectively. Depending on the input transcript, corresponding fluorescence was correctly expressed. Of note, both GFP and BFP were expressed in single cell when two different ORFS transcripts were transfected simultaneously. White arrows indicate cells positive for GFP, BFP, or both.

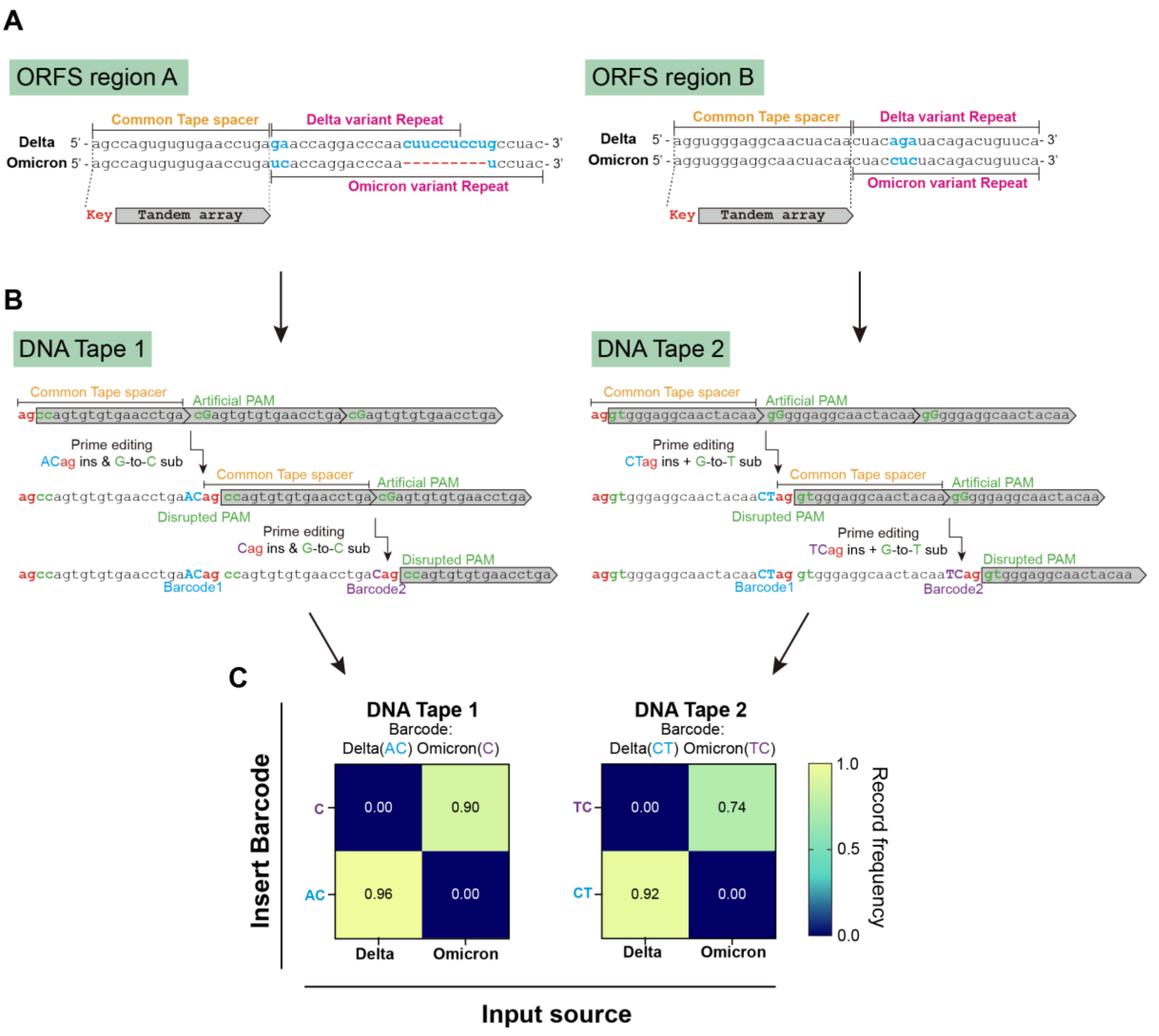


**Fig. S11. Design of two DNA Tapes and unigram recording efficacies of each DNA Tape using CHEETAH with peuRptr-v2s and NG-PE7. (A)** Alignment of two different ORF transcripts (Delta and Omicron) within the selected regions for designing DNA Tapes. Red dashes indicate deletions, and blue letters indicate mismatches. For recording of two SARS-CoV-2 variants on DNA Tape, the shared sequences (Common Tape space) were selected to enable recording on pRecord-Tape and different sequences were selected for hybridization with variant-specific peuRptr-v2s. **(B)** Diagram of two successive SARS-CoV-2 variants-specific RNA recording events on two DNA Tapes. The DNA Tape consists of a tandem repeat that has a 5'end truncated sequence (grey boxes). The desired prime editing activates the next repeat. For activation of next repeat, a 2-bp barcode and a 2-bp key sequence insertion along with 1-bp substitution (disrupting the PAM sequence) are required. **(C)** The recording frequencies of unigram pattern in position 1 across all designed DNA Tapes. The record frequency represents unigram barcode frequency in position 1 within the total edited reads ($n$ = 3).
